# Side-by-side systematic characterization of novel FPs in budding yeast

**DOI:** 10.1101/2025.06.27.661975

**Authors:** Dennis Botman, Daan H. de Groot, Martijn Wehrens, Kelly van Rossum, Sietse Yska, Brian van der Kieft, Vincent Albert, Noémie Minghetti, Joachim Goedhart, Bas Teusink

**Affiliations:** Systems Biology Lab, AIMMS/ALIFE, Vrije Universiteit Amsterdam, 1081 HV Amsterdam, the Netherlands; Biozentrum and Swiss Institute of Bioinformatics, University of Basel, Basel 4056, Switzerland; Section of Molecular Cytology and van Leeuwenhoek Centre for Advanced Microscopy, Swammerdam Institute for Life Sciences, University of Amsterdam, Amsterdam, the Netherlands

## Abstract

Fluorescent proteins (FPs) have become indispensable for biological research. Since the discovery of the first FP, Aequorea victoria green fluorescent protein (avGFP), new fluorescent proteins are continuously being developed. To achieve optimal experimental results, selecting an FP based on specific characteristics—such as brightness, photostability, photochromicity, monomericity, pH robustness, and fluorescence lifetime—is essential. However, a thorough side-by-side comparison of these characteristics is missing for the latest generation of FPs. Here, we provide a comprehensive characterization in yeast of the most recently developed FPs, including FPs that were codon-optimized for yeast. We believe this provides an excellent compendium for choosing the most suitable FP for research purposes.

## Introduction

Fluorescent proteins are indispensable tools in almost every research field in biology. Even after more than 30 years since the first implementation of GFP, new fluorescent proteins continue to be developed to offer improved or altered features. These features include brightness (either the theoretical brightness-quantum yield times the extinction coefficient - or the practical brightness, which is also determined by cellular factors and the amount of matured fluorescent proteins in a cell), photostability (the ability of an FP to remain fluorescent upon exposure to its excitation light), pH-sensitivity, photochromicity (the behavior to photoswitch to an altered color state or a reversible dark state upon exposure to a high-energy wavelength), monomericity (the tendency of FPs to aggregate into oligomers) and fluorescence lifetime (the average time a fluorescent molecule spends in its excited state before emitting a photon and returning to its ground state). Lastly, FP codon-optimization for the organisms of choice is often performed, although its effect on FP performance is not fully understood^1^ and more information on this is desired.

To select the most suitable FP for an experiment from the wide variety of conventional and novel FPs, a systematic characterization in, preferably, identical circumstances is desired. In the recent years, various characterizations have been performed *in vitro*^2^, in mammalian systems at 37°C^3^, or in other organisms growing at lower temperatures^1,4,5^. Yet, a full side-by-side characterization of novel FPs and the current standard FPs is currently lacking. Therefore, in continuation of our previous report^1^, we characterized a wide variety of FPs *in vivo* using budding yeast as model organism growing at 30°C. We included almost all novel FPs developed in the past few years and made codon-optimized variants for Baker’s yeast of the best performers^6–27^. This provides a compendium of the most promising FPs that can aid researchers in choosing the most suitable FP for their experiments.

## Material and Methods

### Generation of 2A constructs

ymTq2-P2A-ymScarletI, ymTq2-O2A-ymScarletI, ymTq2-E2A-ymScarletI, ymTq2-T2A-ymScarletI and ymTq2-ymScarletI were constructed by performing a PCR on a plasmid containing ymTq2 (Addgene plasmid 118453) using the primers listed in table 1. Next, the products and the plasmid containing ymScarletI in pDRF1-GW (Addgene plasmid 118452) were digested using XmaI and NheI (New England Biolabs, Ipswich, MA, USA), whereafter the products were ligated using T4 DNA ligase (New England Biolabs).

**Table 1.**
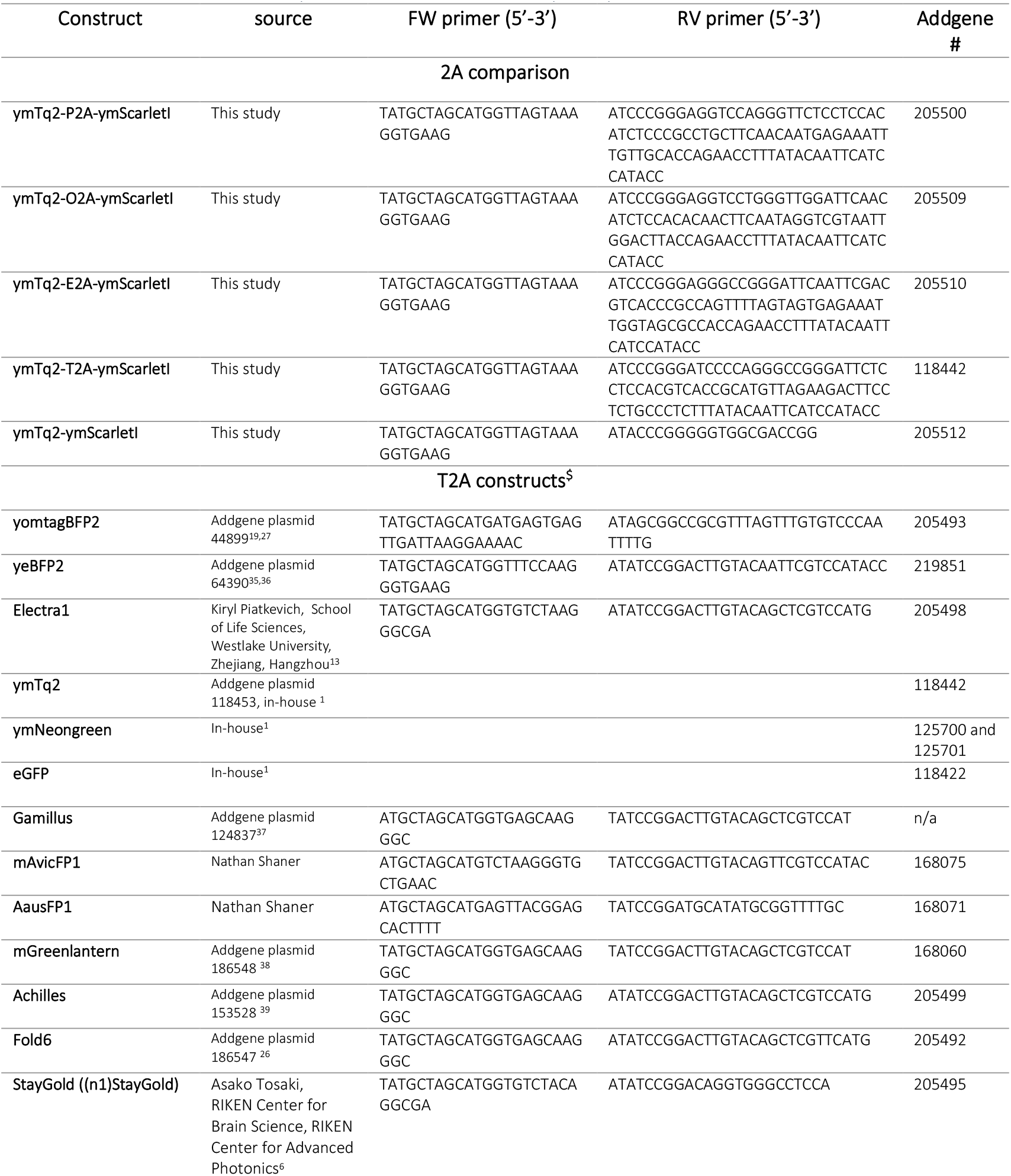
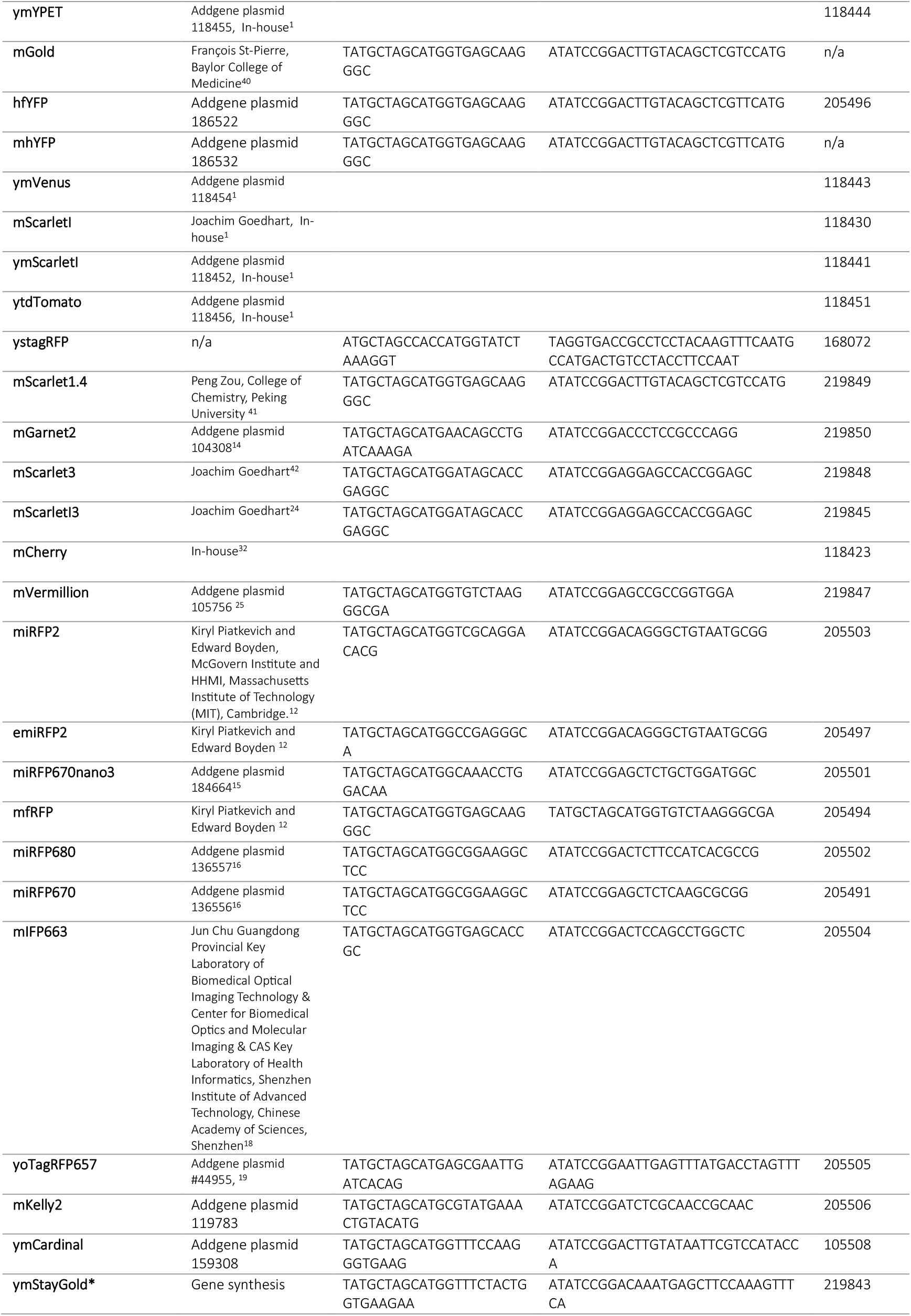
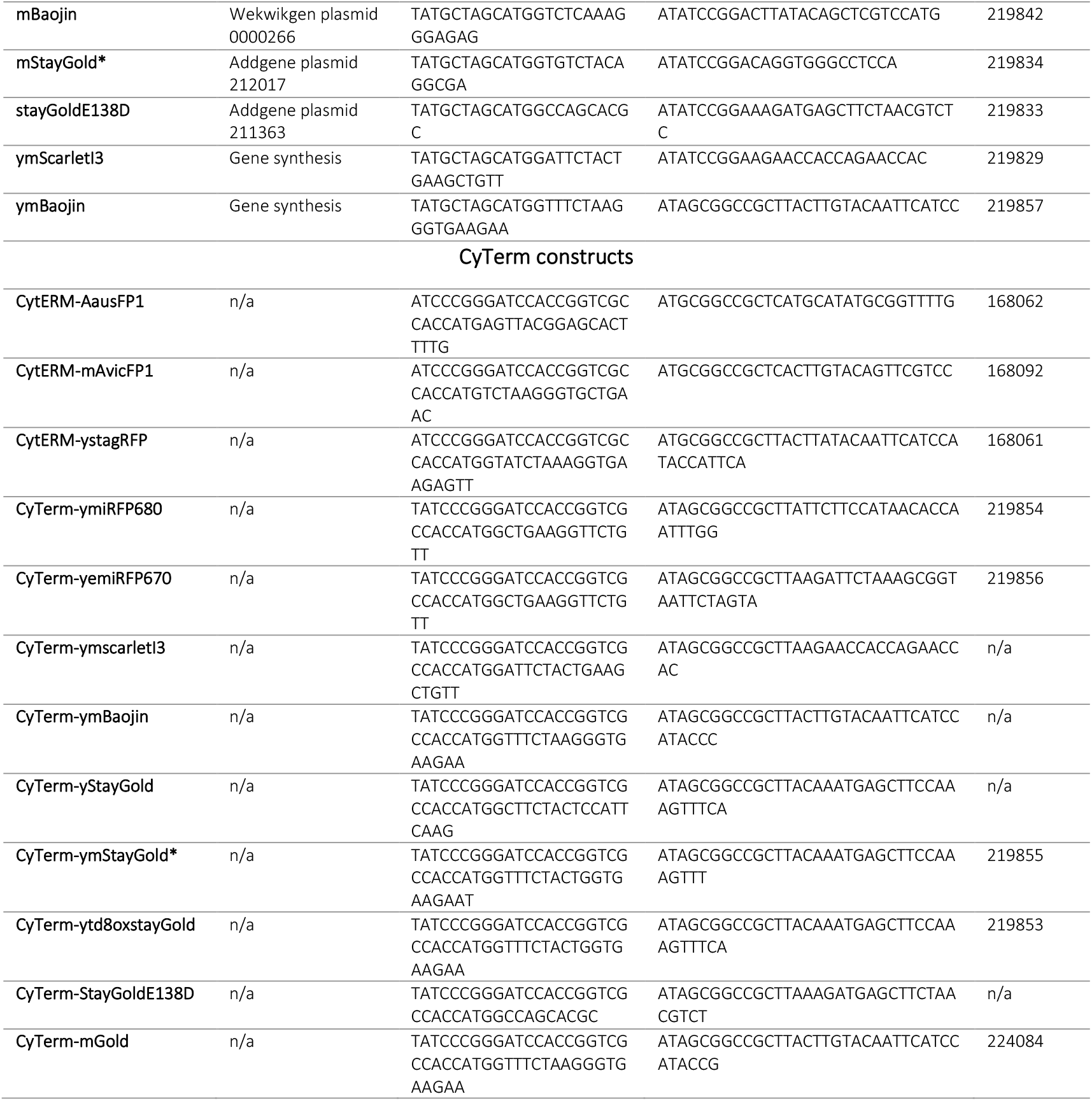
All PCR primers used in this study, including the source of the FPs and the Addgene plasmid number if available. *ymStayGold is the variant published by Ando and colleagues^*8*^. ^$^BFPs, CFPs and GFPs are fused to T2A-mCherry where YFPS, RFPs and iRFPs are fused to T2A-mTurquoise2. All constructs are in the pDRF1 plasmid.

### Generation of T2A constructs

All FPs except ystagRFP and miRFP670 were amplified using KOD one polymerase (Sigma-Aldrich, St. Louis, Missouri, USA) and the primers listed in table 1. Next, all products together with a plasmid containing either FP-T2A-mTq2 or FP-T2A-mCherry in pDRF1 were digested using NheI and BspeI (Thermo Fisher Scientific). Afterwards, the PCR products were ligated in the T2A-mTq2 (for all iRFPs, RFPs and YFPs) or T2A-mCherry (for all BFPs, CFPs and GFPs) plasmids using T4 DNA ligase which generated the T2A constructs. To generate ystagRFP-T2A-mTq2 in pDRF1, a PCR was performed using KOD one polymerase and the primers depicted in table 1. Afterwards, the product and yotagRFPT-T2A-mTq2 (Addgene plasmid #168089) were digested using NheI and BsteII (New England Biolabs) to replace the first part of yotagRFPT with the PCR product containing the mutation to ystagRFP. miRFP670 was generated by performing a PCR on miRFP670 using the primers in table 1 and KOD one polymerase. Afterwards, the product and tdTomato-T2A-mTq2 in pDRF1 were digested with NheI and BspeI and the 2 parts of miRFP670 were ligated into the plasmid generating miRFP670-T2A-mTq2 in pDRF1. yemiRFP670, ymiRFP680, yStayGold and ytd8oxSG were synthesized and digested using NheI and BspeI and ligated in either FP-T2A-mTq2 or FP-T2A-mCherry in pDRF1 digested with the same enzymes which generated the FPs with T2A-mTq2 (for yellow, red and infrared FPs) or T2A-mCherry (for cyan and green FPs).

### Generation of Cyterm constructs

For all CytERM constructs except CytERM-miRFP680, plasmids containing the desired FPs were amplified using KOD one polymerase and the primers listed in table 1. Next, all products and CytERM-mTq2 in pDRF1 were digested using SacII and NotI (for Gamillus) or XmaI and NotI (for all other products) and the products were ligated (using T4 DNA ligase) in the plasmid, replacing mTq2 for the FP of interest. CytERM-miRFP680 was made by performing a KOD one PCR on CytERM-mTq2 in pDRF1 using 5’-agcggccgcACC-3’ and 5’-GGTGGCGACCGGT-3’ as FW and RV primer, respectively. In addition, miRFP680 was amplified using KOD one and the primers listed in table 1. Next, a Gibson assembly (New England Biolabs) was performed on the 2 PCR products, generating CytERM-miRFP680 in pDRF1. Other CytERM constructs were already in-house from a previous publication^1^ and can be found on Addgene as well.

### Yeast transformation

W303-1A WT (MATa, leu2-3/112, ura3-1, trp1-1, his3-11/15, ade2-1, can1-100) cells were scraped from an agar plate, resuspended in a transformation mixture (240 μL PEG 3350 50% w/v, 40 μL 1M lithium acetate, 10 μL UltraPure™ Salmon Sperm DNA Solution 10.0 mg/mL (ThermoFisher Scientific) and 500-1000 ng of plasmid DNA) and incubated for 15 minutes at 42°C. Next, cells were spun down (15000 g for 30 seconds) after which the supernatant was replaced by sterile water. Finally, cells were plated out on selective plates.

### Brightness, photostability and photochromism assessment

W303-1A wild-type (WT) cells expressing the T2A constructs were grown overnight at 200 rpm and 30 °C in YNB medium (Sigma Aldrich, St. Louis, MO, USA), containing 100 mM glucose (Boom BV, Meppel, Netherlands), 20 mg/L adenine hemisulfate (Sigma-Aldrich), 20 mg/L L-tryptophan (Sigma-Aldrich), 20 mg/L L-histidine (Sigma Aldrich) and 60 mg/L L-leucine (SERVA Electrophoresis GmbH, Heidelberg, Germany). The next day, cultures were diluted and grown again overnight to an OD_600_ of 0.2-1.5. Next, cells were transferred to a glass slide and visualized at 30 °C using a Nikon Ti-eclipse widefield fluorescence microscope (Nikon, Minato, Tokio, Japan) equipped with an Andor Zyla 5.5 sCMOS Camera (Andor, Belfast, Northern Ireland) and a SOLA Lumencor light engine (Lumencor, Beaverton, OR, USA). CFPs were imaged using a 438/24 nm excitation filter, a 458LP dichroic mirror and a 483/32 nm emission filter. GFPs were imaged using a 480/40 nm excitation filter, a 505LP dichroic mirror and a 535/50 nm emission filter. YFPs were visualized using a 500/24 nm excitation filter, a 520LP dichroic mirror and a 542/27 nm emission filter. RFPs were visualized by a 570/20 nm excitation filter, a 600LP dichroic mirror and a 610LP emission filter. iRFPs were excited by a 630/38 nm excitation filter, passed through a 655LP dichroic mirror and detected at 698/70 nm (all filters from Semrock, Lake Forest, IL, USA). A 60x plan Apo objective (numerical aperture 0.95) was used. Images were recorded using 80% light power, 2×2 binning and 100 msec exposure time. Bleaching data were recorded by imaging only the channel of interest every 300 msec for 75 seconds. Photochromism data were collected by imaging the FP of interest and the CFP channel every 3 seconds for 75 seconds. Data were analyzed with FiJi (ImageJ version 1.54f, NIH, Bethesda, MD, USA) using an in-house macro that performs the following steps: If needed, drift correction was applied using the image stabilizer plugin^28^. Next, background subtraction was performed using the Fiji background subtraction function with a rolling ball radius of 100 pixels. Afterwards, cells were segmented using the Weka segmentation plugin v3.3.4^29^. The resultant segmented image was made binary, whereafter the “Open”, “Fill Holes” and “Watershed” function is applied. Finally, ROIs were detected using the “Analyze Particles” function. For brightness assessment, the fluorescence of the FP of interest was divided by the fluorescence of the reference FP. Day-to-day variation of each FP is determined by calculating the standard deviation of the mean fluorescence per day whereafter this SD is divided by the mean brightness value of the FP to get the normalized day-to-day variation (i.e. the coefficient of variation). All bleaching curves were fitted for a onephase (equation 1) and a twophase (equation 2) decay curve after which the best fit were chosen based on the Bayesian Information Criterion (BIC) value. Furthermore, the frame number where each FP reached an absolute 50% loss in signal and a relative 50% loss (i.e. between the starting fluorescence and the end fluorescence) was determined.

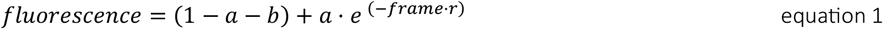

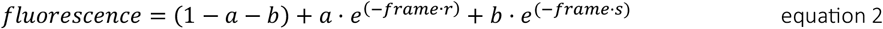

Photochromicity was assessed as described before^1^. In short, to assess whether the FPs can show reversible bleaching under exposure to different wavelengths, i.e., photochromicity, the single and dual excitation curves were modelled using linear differential equations that describe the abundance of 3 FP-states: a fluorescent (natural) state, a bleached state, and an irreversible-bleached state (see our previous publication for more details^1^). The fitted parameters were used for identifying photochromic FPs.

### pH sensitivity assessment

All FPs in the study were evaluated for pH sensitivity using largely the same method as before^1^. The cells were grown overnight in YNB medium as described in “Brightness, photostability and photochromism assessment” to stationary phase (OD_600_ 2.5-3.0). Next, cells were centrifuged at 3000 g for three minutes and the supernatant was removed to concentrate the OD_600_ ten times. Next, a 96 well-plate was filled with 176 μL of a citrate-phosphate buffer (0.1 M citric acid (Sigma Aldrich), 0.2 M-Na_2_HPO_4_ (Sigma Aldrich)) set at pH values ranging from 3 to 8. Next, 4 μL of 2,4-dinitrophenol (DNP, Sigma Aldrich) was added to a final concentration of 2 mM to equilibrate the intracellular pH with that of the buffer^1,30^. Finally, 20 μL of the cells was added and incubated for one to two hours at 30 °C. Fluorescence was recorded using a Clariostar or Clariostar Plus plate reader (BMG labtech, Ortenberg, Germany) at 30 °C. BFPs were measured using 380/15 nm excitation, 403.8LP dichroic mirror and 440/40 nm emission. CFPs were measured using 430/15 nm excitation, 453.8LP dichroic mirror and 480/20 nm emission. GFPs were measured using 470/20 nm excitation, 492.5LP dichroic mirror and 522.5/35 nm emission. YFPs were measured using 497/20 nm excitation, 519.8LP dichroic mirror and 550/35 nm emission. RFPs were measured using 544/15 nm excitation, 565.8LP dichroic mirror and 600/40 nm emission. iRFPs were visualized using either 585/70 or 630/60 nm excitation with a 634LP or 675LP dichroic mirror and 675/54 nm or 715/50 nm emission. Fluorescence signals were corrected for autofluorescence by subtracting fluorescence from cells expressing an empty pDRF1 plasmid and fluorescence was normalized to the pH giving the highest fluorescent signal. Fluorescence values that became negative after background subtractions were discarded. For some FPs, only the codon-optimized or the non-codon optimized variant were assessed as codon optimization does not affect pH curves (F-test, p > 0.05). Next, all curves were fitted against a Hill fit (equation 3 with b being the bottom fluorescence signal, and n the Hill coefficient) or a Gaussian fit (equation 4), where pH_optimum_ indicates the pH at which the fluorescence is maximal, and se^2^ describes the rate at which the fluorescence drops when the pH is suboptimal.

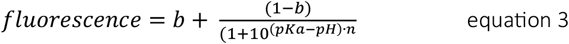

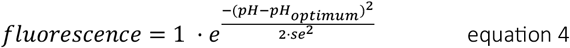

### Assessment of monomericity

The best-performing FPs regarding brightness, photostability and photochromicity were tested for monomericity using the OSER assay^2,31^. As non-monomeric reference FPs, dTomato and yeVenus were used^2,32^, (y)mTurquoise2 was included as monomeric reference FP^1,2,33^. The strains expressing the CyTerm-FP constructs were grown overnight in YNB medium as described in the “Brightness, photostability and photochromism assessment” section to stationary phase (OD_600_ 2.5-3.0), incubated for at least 1 hour at 18-20 °C and placed on a glass slide and visualized with a Plan Apo λ 100x Oil Ph3 objective (numerical aperture 1.45) and the same filter settings as described in “Brightness, photostability and photochromism assessment”. Next, a z-stack of 9 steps, 0.5 μm per step, was captured using light power and exposure times that gave sufficient signal. Afterwards, a MAX project of slides 3-7 was made and an in-house macro which uses the random-forest WEKA segmentation plugin v3.3.4 of Fiji was used to identify OSER structures^1,29,34^. The number of cells without any OSER structures is the OSER score and hence a value for monomericity of a certain FP.

### Fluorescent lifetimes

W303-1A WT cells expressing either the T2A constructs or a single FP were spotted on a 2% agarose plate containing the same composition as the media described in the “Brightness, photostability and photochromism assessment” section. Next, the plates were incubated for at least three days at room temperature and whole colonies were imaged to determine lifetimes using a Leica Stellaris 8 equipped with a pulsed white light laser set to 40 MHz and a 1.25x NA 0.04 objective. Lifetimes of FPs of different color classes were measured using the following settings: for cyan excitation at 440 nm and detection between 450-502 nm on a HyD X 2 detector, for yellow and green excitation at 488 nm and detection between 502-560 nm on a HyD S 3 detector, for red excitation at 570 nm and detection between 580-649 nm on a HyD S 3 detector, and for far red excitation at 642 nm and detection between 651-755 nm on a HyD R 5 detector. Laser power was set to 3%, the pinhole to 2 airy units, the imaging speed to 100 Hz per line with 3 line repetitions and a frame size of 512×512.

### Sampling details & data analysis

For all microscopy data, all cells picked up in the segmentation analysis were analyzed. For the brightness analysis, cells that have a fluorescence of approximately 2.5 times lower than the background in either the CFP or the RFP channel were discarded. Also, cells with a mean value above 60,000 (arbitrary units) were discarded as these are saturated. Lastly, cells having an FP brightness >15 compared to the reference FP were excluded, as these are typically found to consist of debris or dead cells. For bleaching curves, cells that have a frame with a mean value above 58,000 (arbitrary units) or below 100 (arbitrary units) were discarded. For photochromicity curves, cells that have a frame with a mean value above 60,000 (arbitrary units) or below 100 (arbitrary units) were discarded. For pH curves, fluorescence values that became negative after background subtractions were discarded. For monomericity assessments, no data points were excluded. For lifetime experiments, samples with a low fluorescence signal compared to background (approximately <3x) were discarded.

## Results

### FP brightness

Two properties of FPs that are commonly evaluated to choose a suitable FP are the brightness and photostability. Here, we assessed *in vivo* brightness in budding yeast using the polycistronic T2A element which generates unimolecular concentrations of the FP of interest together with a second FP that serves as a reference to normalize the fluorescence value (figure 1). We tested which 2A sequence was most efficient for separation of the 2 FPs by measuring acceptor emission of ymTq2-[2A]-ymScarletI FRET pairs (fig. S1). No significant differences between the 2A variants were found, which made us decide to continue with the T2A sequence used before^1^.

**Figure 1.**
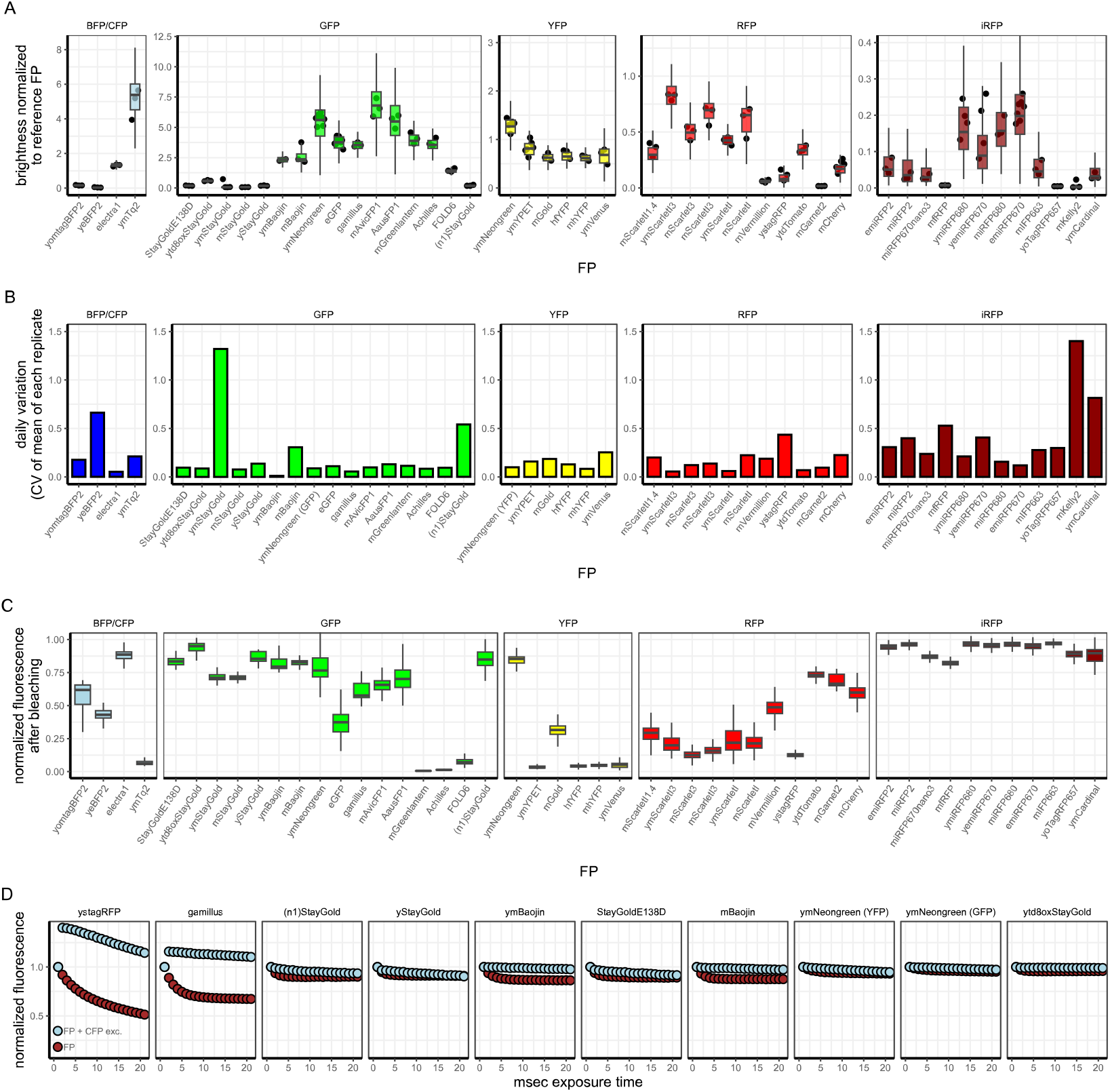
Brightness, day-to-day variation, photostability and photochromicity assessment of FPs. A) fluorescence brightness of FPs compared to either mTq2 or mCherry. Each point depicts the median brightness of all cells measured of a biological (i.e. daily) replicate. Boxes indicate median with quartiles based on the medians of all biological replicates; whiskers indicate largest and smallest observations at 1.5 times the interquartile range. Spectral class is depicted above each facet. B) Daily variation of all FPs. Day-to-day variation is depicted as the coefficient of variation between the mean brightnesses of all daily replicates per FP. Each facet depicts a spectral class (i.e. FP colour), n ≥ 160 and from at least 3 different days for each FP. C) Final fluorescence normalized to its initial fluorescence depicted in boxes for each FP. Boxes indicate medians with quartiles; whiskers indicate largest and smallest observations at 1.5 times the interquartile range. Spectral class (i.e. FP colour) is depicted above each facet, n ≥ 23 cells and from at least 3 different days for each FP. D) Bleaching traces of the identified photochromic FPs indeed show altered bleaching kinetics when co-excited with 426-450 nm. Dots show fluorescence values, normalized to the first frame. Colors indicate excitation setting, n ≥ 76 cells from at least 3 different days.

We found significant differences in brightness between FPs, also within the spectral colour groups itself (ANOVA, p<0.01 and Tukey multiple pairwise-comparisons). For BFPs and CFPs, we found ymTurquoise2 (ymTq2) was the brightest FP. However, yomtagBFP2 and yeBFP2 are not optimally visualized using our CFP filter set. These FPs might still be useful when a truly blue FP is needed. When a CFP is needed, ymTq2 is still the preferred protein in terms of brightness. For GFPs, we found that mAvicFP1 and AausFP1 are bright FPs and potentially perform even better than ymNeongreen, especially since we found that codon-optimalisation can improve *in vivo* brightness for GFPs^1^. All StayGold variants show a remarkably low brightness where (y)mBaojin is the only one that stood out and has an acceptable brightness. For the YFPs, ymNeongreen is still the best performer (although not a pure YFP) and ymYPET the brightest pure YFP. ymScarletI3 is the brightest RFP among all new FPs tested. Finally, iRFPs are rather dim using our setup, where miRFP680 and emiRFP670 stand out and are brighter compared to the other iRFPs. Furthermore, we determined how much variation in brightness exists between daily replicates for each FP. To do so, we determined the mean brightness of biological replicates for each day for each FP and determined the coefficient of variation over these values. Here we found that most FPs have a rather comparable daily variation. Only yeBFP2, (n1)StayGold, mfRFP, mKelly2 and ymCardinal showed a notably higher coefficient of variation. This should be considered when performing quantitative measurements between different days using these FPs.

Even a bright FP is impractical to use if its signal diminishes too fast. Thus, we assessed photostability of all FPs by making a timelapse of cells expressing the FPs and measuring the fluorescence decrease over time (figs. 2 and S2). Next, a one-phase or two-phase decay curve was fitted and the fitted parameters are accessible in table S1. However, some FPs do not reach an absolute 50% bleaching and often reach a plateau above 50% with our settings (which are standard imaging settings). Fits to such curves also tend towards a plateau above 50% and this biases the t_1/2_ values. In other words, photostable FPs can have a low t_1/2_ value if they hardly bleach. Therefore, we prefer to depict the final fluorescent fraction at the end of the bleaching period (fig. 1C). Photostability values were different between FPs (ANOVA, p<0.01). For CFPs, we found that ymTurquoise2 (the brightest CFP) is not photostable, but Electra1 is. For GFPs, all StayGold variants show high photostability as expected and ymNeongreen, Gamillus, mAvicFP1 and AausFP1 are also photostable. All other GFPs show a low photostability. YFPs, known for their low photostability, show indeed low photostability except for ymNeongreen. If a true YFP is needed, the choice would be mGold. RFPs also show a rather low photostability but ytdTomato, mGarnet2 and mCherry are photostable options. Using our setup, iRFPs show a remarkable high photostability, only miRFP670nano3 and mfRFP show little bleaching while the other hardly show any reduction in fluorescence.

**Figure 2.**
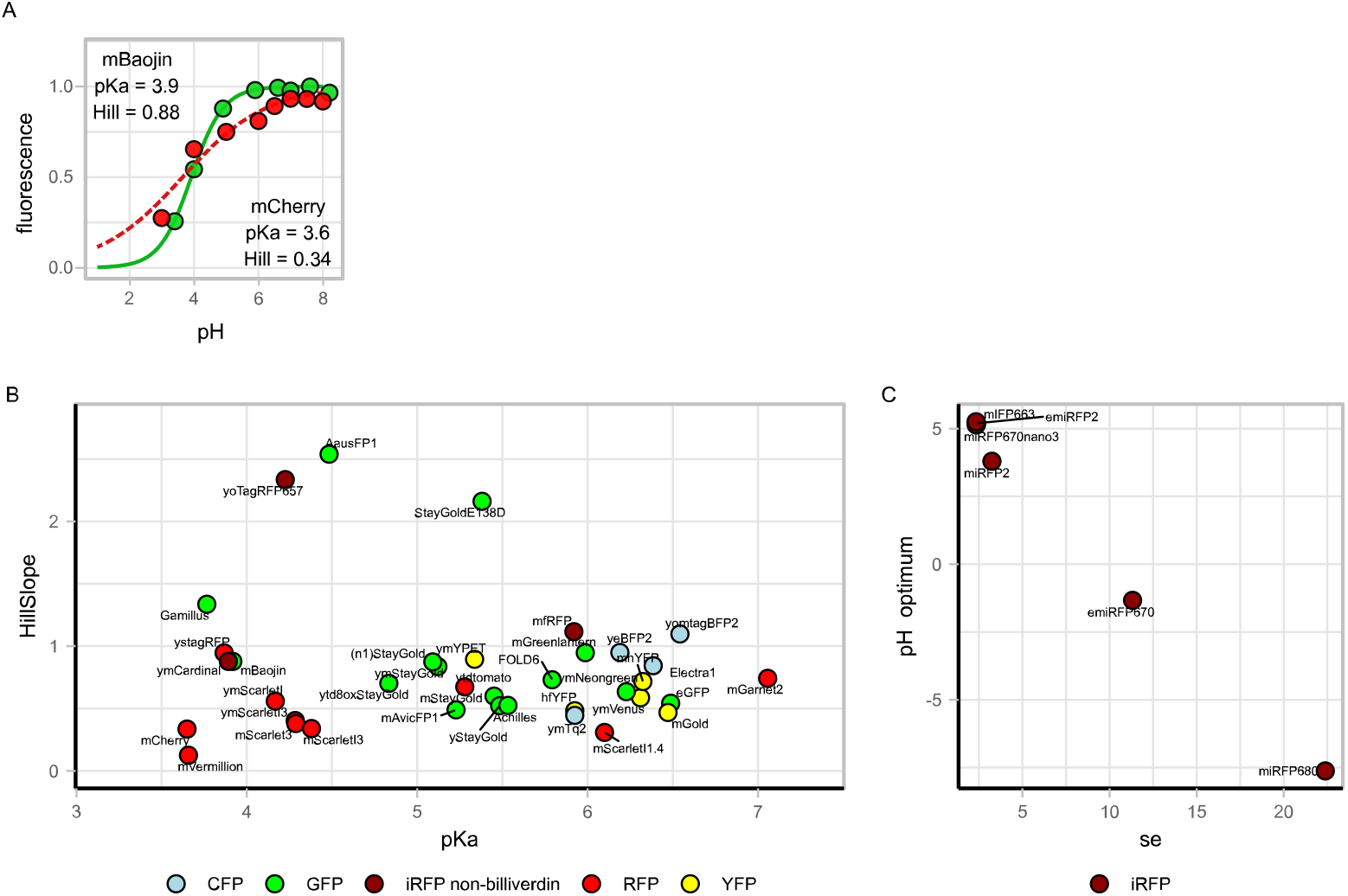
pH sensitivity of FPs. A) Giving only the pKa obscures the actual pH sensitivity. Two pH curves, mBaojin (solid green line with green dots) and mCherry (dashed red line with red dots), are shown. mBaojin has a higher pKa, indicative for a high pH sensitivity, compared to mCherry. However, its Hill coefficient is higher which actually results in a lower pH sensitivity (i.e. the plateau at high pH is reached faster). Lines show the Hill fit, dots show the mean fluorescence value, normalized to the pH giving the highest fluorescence. B) Hillslope and pKa obtained from the Hill fit plotted for each FP. Colour indicates FP colour class. C) se and pH optimum obtained from a Gaussian fit plotted for each iRFP with biliverdin as chromophore. Colour indicate FP colour class. Data is from at least 3 technical replicates per FP.

**Figure 3.**
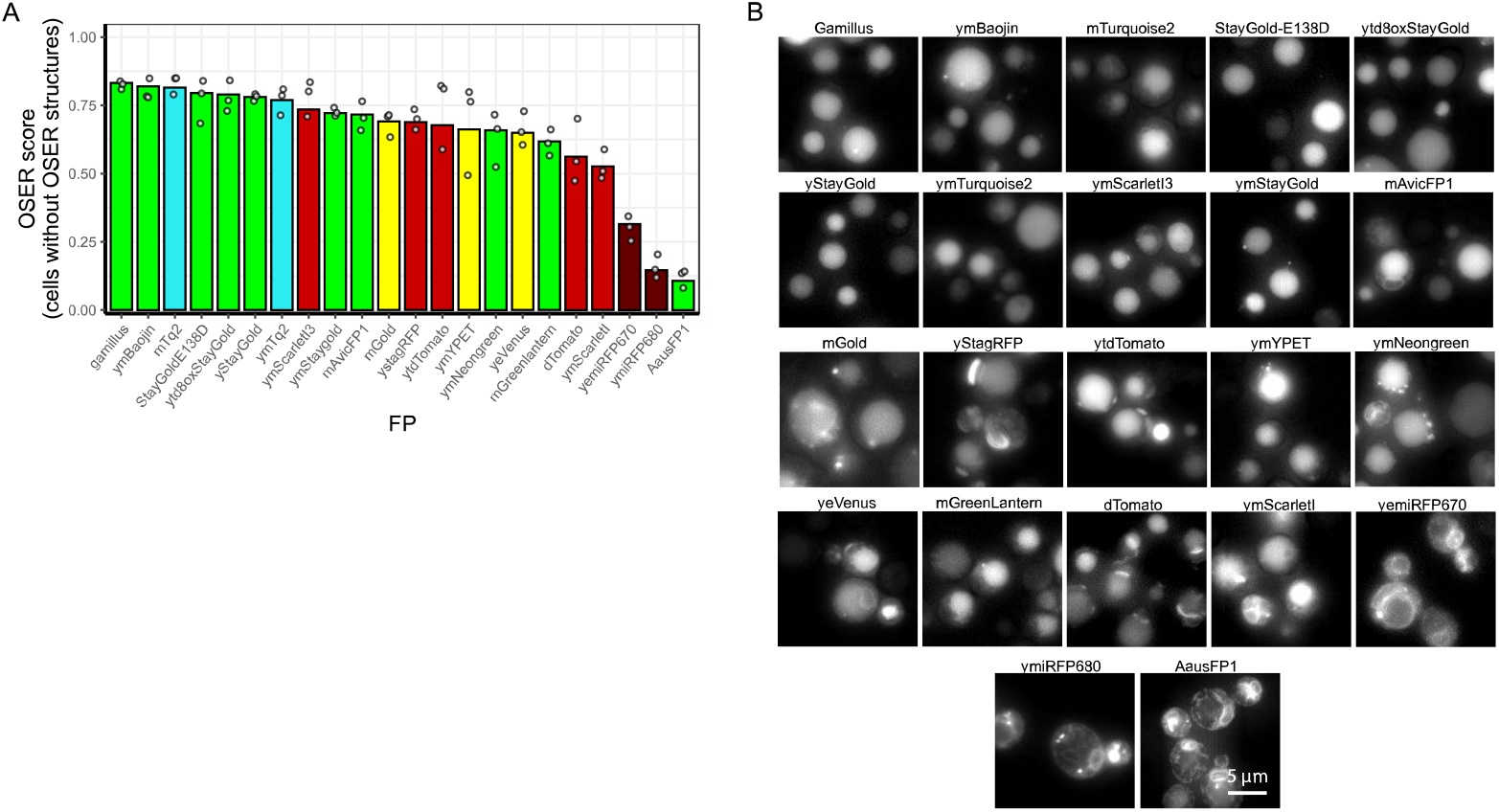
Monomeric properties of 22 FPs. A z-stack was captured of W303-1A cells expressing Cyterm-FP whereafter a Z projection was performed and OSER structures were identified using the random-forest WEKA segmentation plugin of FiJi^*29,34*^. A) OSER score per FP, depicted by the bars. Dots indicate mean OSER score for each biological replicate. Colours indicate FP colour. B) Representative pictures of yeast cells expressing the Cyterm-FP (FP depicted above each picture). Per FP, n ≥ 1370 cells from at least 3 different days.

When FPs are visualized together with a low-wavelength FP (e.g. CFP), reversible bleaching can occur ^43–49^. This happens when FPs can switch between different color states (e.g. fluorescent and non-fluorescent) after exposure to light, a phenomenon called photochromicity. FPs in a non-fluorescent state can be pushed back into the fluorescent state when using short wavelength light. We characterized photochromicity by performing bleaching experiments alternating between the FP and CFP excitation filters. The results were analyzed using our previously published model^1^ and the obtained photochromicity value was used as an indicator of the severity of photochromic behavior (figs. 4 and S3). We found 9 FPs to be photochromic; stagRFP and gamillus show severe photochromicity, followed by (n1)StayGold with modest photochromic behavior and yStayGold, ymBaojin, StayGoldE138D, mNeongreen and td8oxStayGold with minimal photochromic behavior. Of note, we also find photochromic behavior of Electra1, but this FP was not added to this dataset as it is a CFP itself.

**Figure 4.**
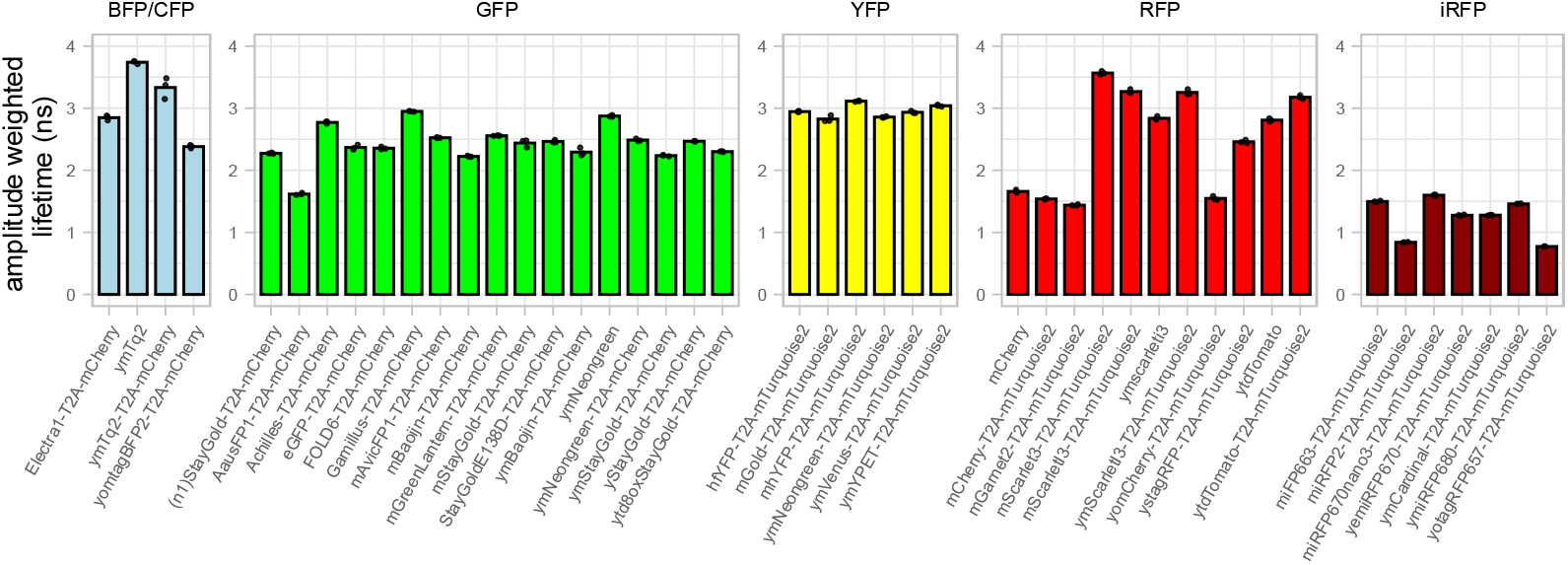
FP lifetimes. W303-1A cells expressing various FPs were spotted on agarose plates and imaged using a Leica Stellaris 8. Lifetimes were obtained using the LAS-X software. Shown are the amplitude weighted lifetimes of at least 3 spotted colonies per FP construct, depicted underneath the bars. Bars show mean lifetimes, spots indicate individual measured colonies. Spectral class is depicted above each facet.

### pH stability

FPs used in an acidic environment or environments that show pH dynamics should exhibit sufficient pH robustness. Although the pKa of many FPs is known, detailed characterization is often lacking. pKa values give an indication of the sensitivity to pH changes, but the hill coefficient (i.e. the steepness of the pH curve) is an important second parameter as this indicates the width of the plateaus above and below the pKa (fig. 2A). A high hill coefficient corresponds to a steep curve through the pKa and results in reaching a stable plateau faster, which is necessary for a pH-robust FP. We recommend to always share these two values together with the pH curve on which these values were estimated.

Using the ionophore 2,4-dinitrophenol to equilibrate intracellular and extracellular pH, we measured fluorescence responses across a range of pH values in yeast cells. The obtained fluorescence values were plotted and fitted with a Hill fit for all FPs with a non-biliverdin chromophore (figs. 2 & S4). For iRFPs that have biliverdin as chromophore, a Hill fit cannot explain the data. A bell-shaped curve is preferred^12,15,18,50^ and we share the parameter (named se) for this curve (indicative of the width of the curve) and the pH optimum for these FPs. For some FPs, we performed a fit only on either the codon-optimized version or the original FP as codon optimization does not change the pH curves (F-test for 5 FPs from Botman et al., 2019^1^, all p-values > 0.05), therefore we refer to all FPs here by their original names. For all non-billiverdin FPs, some stand out in pH robustness. These FPs are AausFP1, TagRFP657, Gamillus, stagRFP, mCardinal, mBaojin, mScarletI, mScarletI3 and mCherry, which all show a low slope in the physiologically relevant pH range of 6.5-7.5 (fig. S4). mVermillion also showed pH robustness in the fit, but the data quality was poor because of the low expression. The most pH-robust YFP is ymYPET and the best CFP is mTurquoise2. For biliverdin-dependent iRFPs, all FPs are robust, with emiRFP670 and miRFP680 being the best performers.

### Monomeric properties

In addition to all characteristics related to fluorescence, FPs should also be monomeric as non-monomeric FPs can affect the function and localization of tagged proteins. The 27 most promising FPs based on the previously characterized properties were assessed for their monomerism using the OSER assay^2,31^. As non-monomeric reference FPs we used dTomato and yeVenus^2,32^. Significant differences in OSER values were found between the FPs (Kruskal-Wallis rank sum test p<0.01). The cyan FP mTq2 showed a high OSER score meaning it is a monomeric FP. For GFPs, mNeongreen, mGreenLantern and AausFP were not monomeric. This is mostly in line with previous results^8,51^ although mGreenLantern was reported to be monomeric^38^. For the YFPs, mGold and —despite some variability— ymYPET showed an acceptable OSER score. Other YFPs were not monomeric. Of the RFPs, ymScarletI3 was monomeric. As expected, yStagRFP and tdTomato are not monomeric. Interestingly, ymScarletI did not exhibit monomeric behavior either, which was less anticipated^1,42^. Lastly, both the bright iRFPs (emiRFP670 and miRFP680) are, in contrast to previous reports^16^, not monomeric in our assay which should be taken into consideration when designing experiments.

### Lifetimes

In recent years, lifetime measurements have become a conventional method to measure FPs since measurement setups have become more accessible and widely available. Therefore, a comprehensive assessment of lifetimes is desired. We measured the lifetimes of almost all assessed FPs in this study by measuring agarose plates with spotted yeast cells (fig. 4). Most of the results were in line with the existing data. For CFPs, mTq2 showed the highest lifetime of 3.7 ns (3.4 for the T2A-mCherry variant). For GFPs, we found high lifetimes for Gamillus and ymNeongreen (2.9 ns), whereas AausFP1 exhibited a short lifetime (1.6 ns). YFPs had similar lifetimes of around 3 ns. RFPs showed the highest variation between FPs in lifetimes and mScarlet3 outperforms all other RFPs with a lifetime of 3.6 ns. mGarnet2 and mCherry showed short lifetimes of 1.4 to 1.6 ns. Lastly, iRFPs showed short lifetimes with miRFP670nano3 having the longest lifetime of 1.6 ns and yotagRFP657 and miRFP2 with the shortest lifetimes recorded among all FPs (0.8 ns). These data can be used to identify candidates for lifetime sensors^52–54^ or for lifetime-based signal unmixing^42,55,56^.

## Discussion

New FPs are constantly being developed to extend and improve the toolbox of single-cell biology. To use the most suitable FP for a certain experiment, a side-by-side full comparison of these novel FPs is necessary. Here we provide such a comparison by assessing *in vivo* brightness, photostability, photochromicity, pH robustness and lifetimes for more than 40 FPs and more than 15 codon-optimized FPs. We found that mTq2 is still the best CFP choice, although Electra1 is more photostable, it is also photochromic. Both FPs are rather pH-sensitive although mTq2 seems slightly more pH robust. For this spectral class, there is room for improvement in terms of photo- and pH-stability. For GFPs, we suggest using mNeongreen or mBaojin. mBaojin and Gamillus were the most pH robust GFPs, but Gamillus showed severe photochromicity (as mentioned by its developers^37^). For YFPs, mNeongreen, mYPET and mGold are bright FPs, where mYPET showed low photostability and proper pH stability. Recently, two new mGold variants, mGold2s and mGold2t, were developed with improved photostability^57^. However, their properties in our experimental setups have yet to be characterized. For RFPs, mScarletI3 and tdTomato were the brightest FPs with tdTomato being more photostable and mScarletI3 being monomeric. pH stability was comparable for these FPs. Here, two new RFPs named mScarlet3-H (or mYongHong) and mScarlet3-S2 were published with improved protein stability and photostability^58,59^. However, these FPs are shown to be rather dim but can be considered when a high photostable RPF is needed. Lastly, for iRFPs we recommend to use emiRFP670 and miRFP680, which -in line with previous results^3^-were bright, photostable and pH stable, but -in contrast to previous results-not monomeric. It is important to note that the obtained characteristics are system-dependent and may vary between different experimental setups. An example is the yeBFP2 brightness: this was obtained using a CFP-optimized filter set, resulting in suboptimal visualization conditions for this FP.

We are surprised by the relatively low brightness of all StayGolds and their derivatives. We saw a relative brightness of 0.06-0.25 for all StayGold variants (with ytd8oxStayGold having a brightness of 0.6), while only mBaojin stood out with a brightness of approximately 2.8 relative to mCherry. This is lower than the previously reported brightnesses although Ivorra-Molla and colleagues also showed that the practical brightness is lower than that of mNeongreen in mammalian cells, which we confirm in budding yeast^6,9,10,60^. mBaojin had a slightly lower photostability, but not as severe as Shimozono and colleagues found (20% of StayGold). mStayGold had an even worse performance compared to mBaojin in our assay. For the brightness, FP termini can affect the practical brightness of StayGold (and probably also other FPs)^9,60^. However, we used untagged proteins in yeast and the differences found here might also be conditions specific. As we previously suggested, temperature is an important factor and StayGold is originally optimized for expression in bacteria at 37 degrees^1^. With many model organisms (e.g. budding yeast, zebrafish, *C. elegans*, fruit flies) having lower optimal temperatures, the impact of this parameter on FP behavior should not be underestimated. Thus, while these results are certainly relevant for research at 37°C, they are especially valuable for studies conducted at lower or alternative temperatures.

Our systematic assessment of monomerism of the FPs is largely in line with previously published results^2,6,24^. Yet, three things stood out. We found StayGold to be a monomeric FP which is unexpected as it is published as a dimer. Yet, the original publication already showed an OSER score comparable to mNeongreen, which indicates it is not a strong dimer. Potentially, the organism and experimental conditions of choice determine the readout of the OSER assay or expression levels affect OSER formation (as mentioned by Costantini and colleagues^31^). Secondly, we found that mScarletI was a poor performer in terms of monomerism, but this is solved for mScarletI3. mScarletI3 has new termini and mutations on the outside of the β-barrel which probably improve monomeric properties. Lastly, we found that the iRFPs were far from monomeric and this has to be taken into consideration for experiments. We believe that in this spectral range improvements can be made, both in terms of brightness and monomeric properties. Recently the FluoPPI assay was published as a new method to determine monomeric properties^61^. Although the OSER assay shows highly similar -yet not identical-results compared to this method^8,60^, it can be worthwhile to include the FluoPPI assay as a second control for monomeric properties in the future. Still, the ‘m’ prefix in many FP names has lost its meaning as many mFPs exists that are shown to be far from monomeric (see fig. 7). Our systematic OSER assessment can aid researchers to truly check the monomeric property of their considered FP for an experiment.

Lifetimes of FPs were as expected with mTq2 and mScarlet3 being the highest among all FPs. For GFPs, mNeongreen (in agreement with literature^62^) and Gamillus had rather long lifetimes with little differences found for YFPs. Lastly, in line with previous results^3,63^, iRFPs had short lifetimes likely due to their altered chromophore compared to other FP classes, or due to the inverse relationship between lifetime and fluorescence observed for GFP-based iRFPs^63^.

For the best performing new FPs, we created codon-optimized variants for use in yeast. Previously we found decreased brightness for codon-optimized RFPs^1^, which we attributed to impaired folding. This does not hold for ymScarletI3; the codon-optimized version has improved practical brightness compared to mScarletI3 and perhaps the mutations in this novel FP improves folding and maturation dynamics in yeast.

Here, we do not propose the “best FP”, as it simply does not exist. However, we believe we provide a comprehensive overview, including Table S1 showing all assessed characteristics which should be used to pick the optimal FP for an experiment which hopefully contributes to improved experimental results.

## Acknowledgements

We thank dr. Andreas Milias-Argeitis (Molecular Systems Biology, Groningen Biomolecular Sciences and Biotechnology Institute, University of Groningen, Groningen) for critically reading the manuscript.

## Data availability

All data can be found at 10.5281/zenodo.14711596.

## Supplements

**Figure S1.**
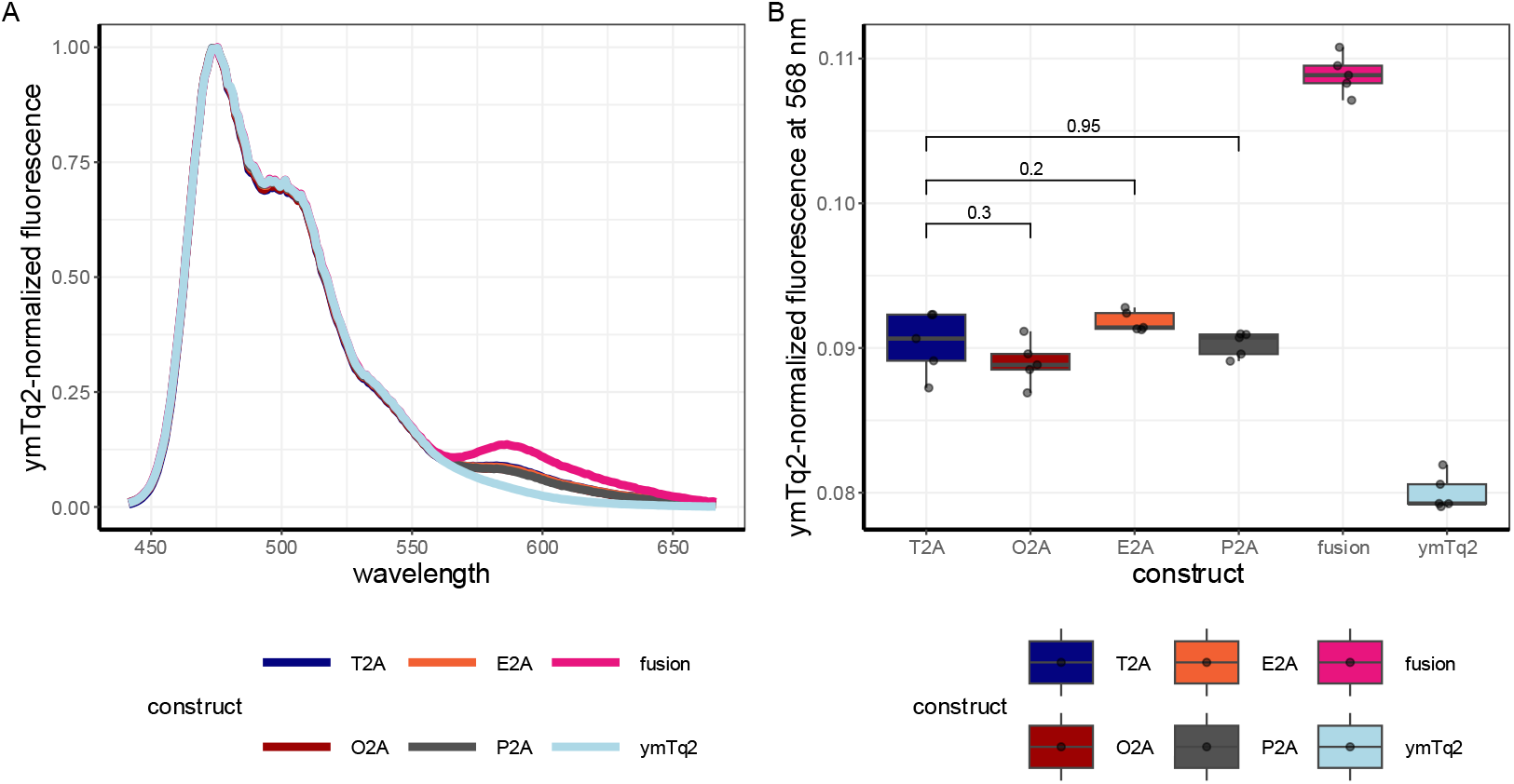
Comparison of 2A variants. A) Emission spectra of W303-1A yeast cells expressing ymTq2, ymTq2-ymScarletI or the four ymTq2-2A-ymScarletI variants. B) comparison of the FRET-induced signal (fluorescence at 568 nm) of the 6 constructs. Values above the whiskers indicate Student’s T-test p value.

**Figure S2.**
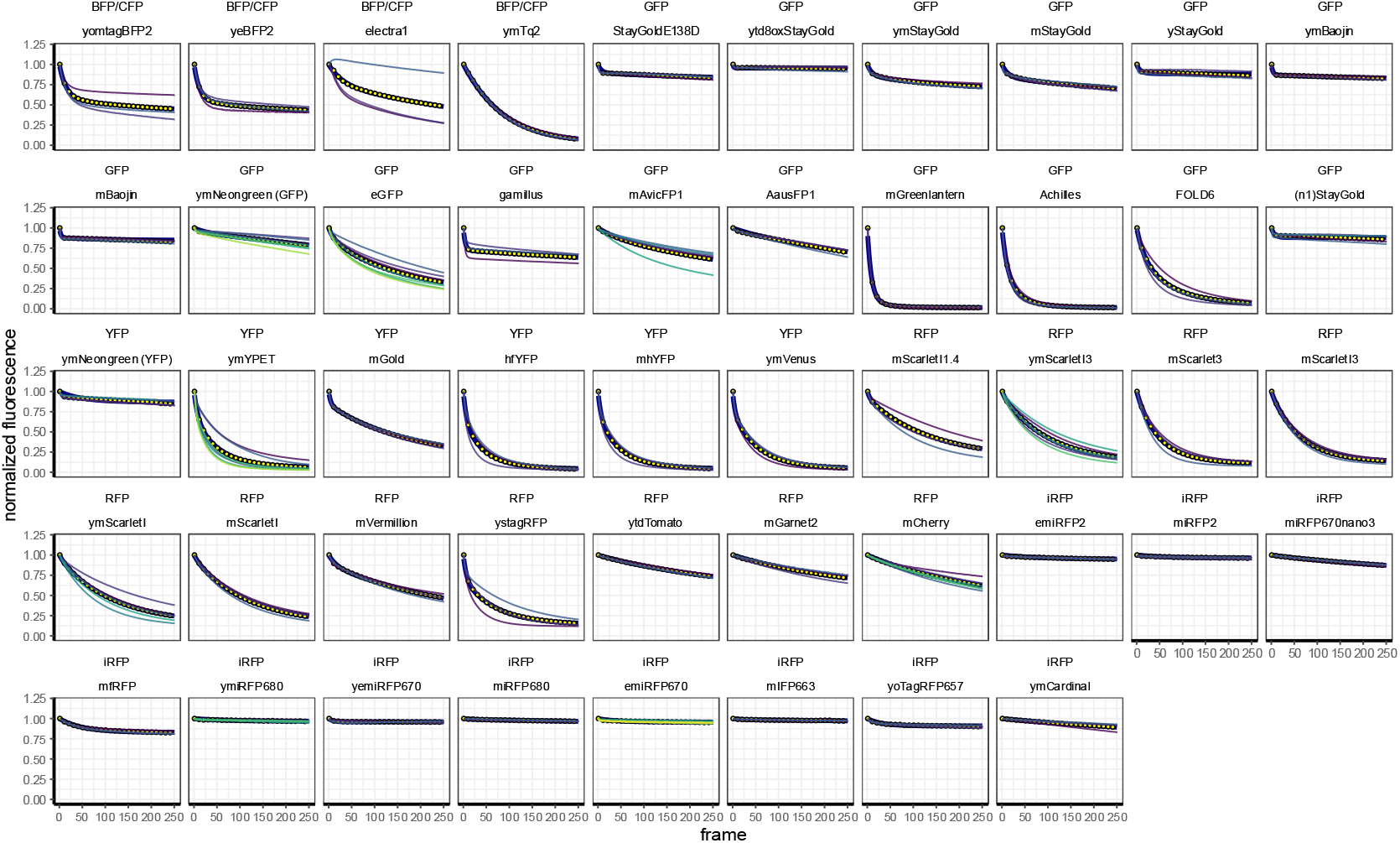
Bleaching curves of all assessed FPs. W303-1A cells expressing the FP-T2A-FP_control_ (depicted above each facet, together with its spectral colour) were visualized for 251 frames and the fluorescence signal was recorded, normalized to the first frame after which a one-phase or two-phase bleaching fit was performed (according to the Bayesian Information Criterion value). Yellow dots show mean fluorescence values, thin lines show biological replicates. Darkblue line shows the fit. Data of at least 3 biological replicates are shown.

**Figure S3.**
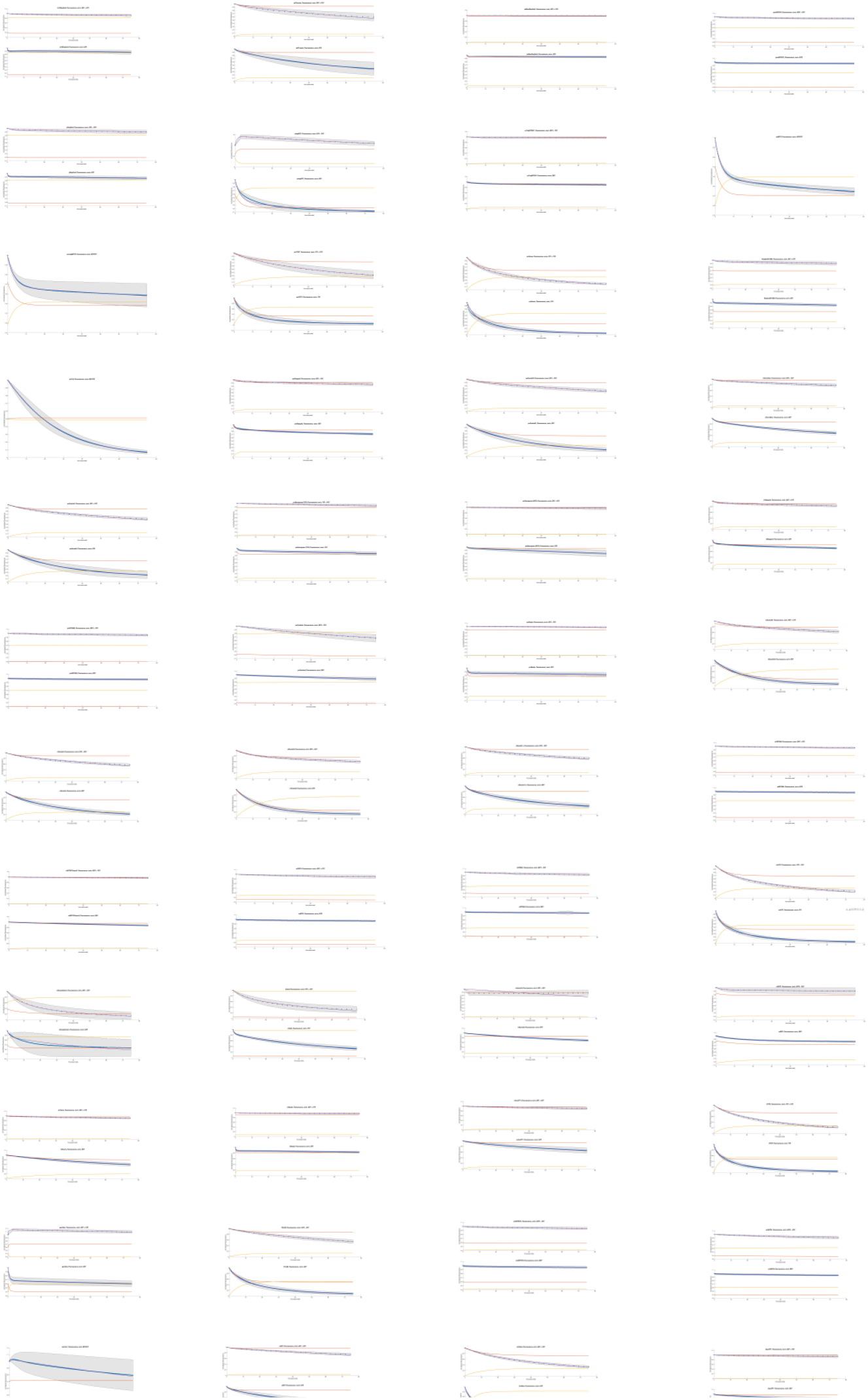
Photochromicity fits of all assessed FPs. W303-1A cells expressing the FP-T2A-FP_control_ were visualized for 26 frames using excitation settings of the FP of interest and CFP excitation (438/24 nm) and the fluorescence signal was recorded, normalized to the first frame after the previously published photochromicity fit was performed^1^. Purple dots and purple shade indicate mean fluorescence values and SD, respectively. Data of at least 3 biological replicates are shown. Red curve indicate the fraction of FPs in the fluorescence (natural) state, yellow the fraction in the reversible dark state and purple the fraction in the fluorescence (natural) state, normalized to its starting value.

**Figure S4.**
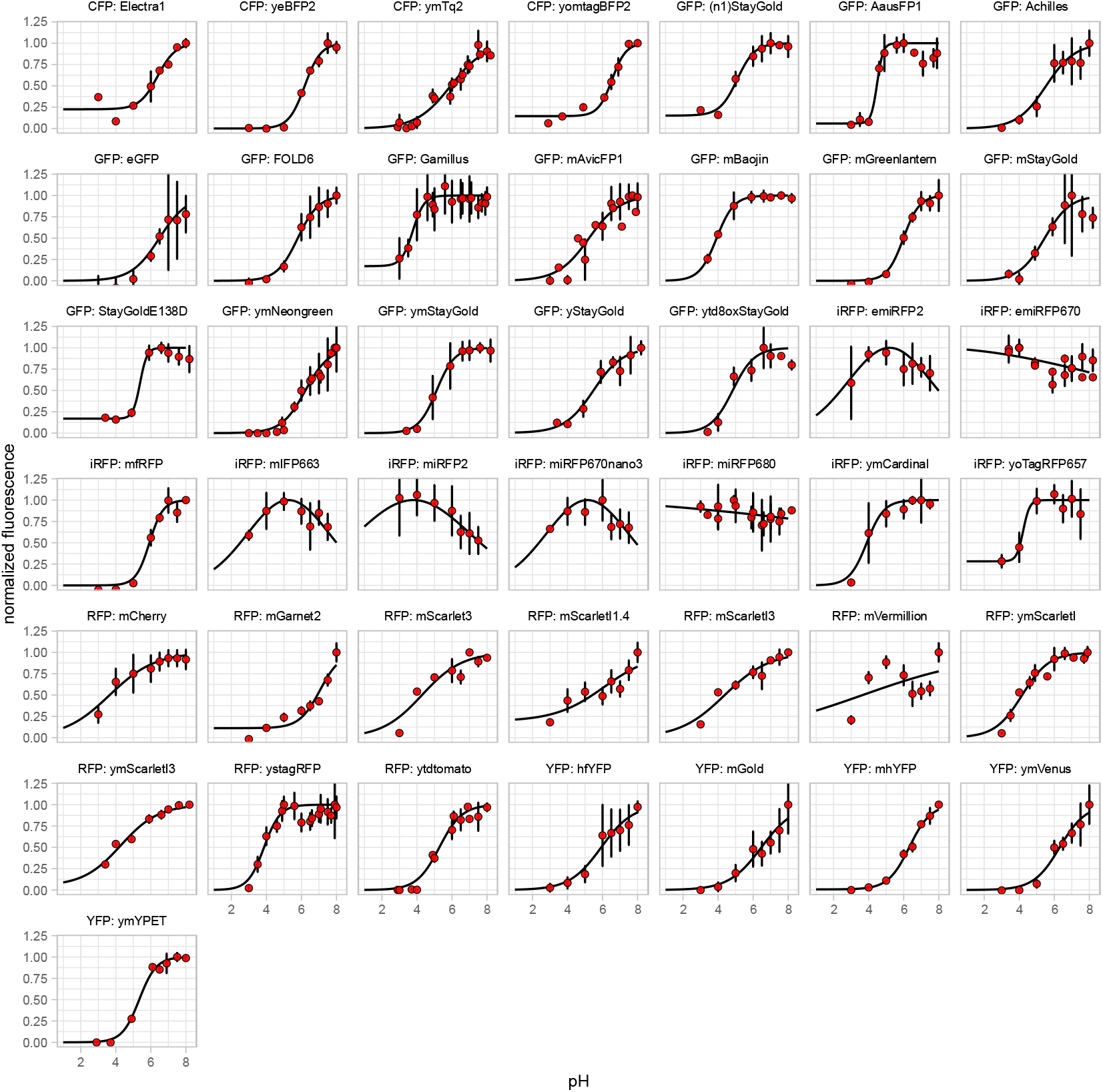
pH curves of all assessed FPs. Cells expressing FP-T2A-FP_control_ were grown overnight, washed twice and resuspended in a citrate phosphate buffer set at pH values ranging from 3 to 8 together with 2 mM 2,4-dinitophenol. Fluorescence was recorded and normalized to the pH value giving the highest fluorescence. Next, a hill-fit or a Gaussion fit was performed for non-biliverdin FPs or biliverdin FPs, respectively. Dots show mean normalized fluorescence, error bars indicate SD. Black line shows the fit. Each facet title depicts the FP and the spectral colour. Data of at least 3 technical replicates are shown.

